# Inhibitory and Stimulatory Micropeptides Preferentially Bind to Different Conformations of the Cardiac Calcium Pump

**DOI:** 10.1101/2021.06.15.448555

**Authors:** Sean R. Cleary, Xuan Fang, Ellen E. Cho, Marsha P. Pribadi, Jaroslava Seflova, Jordan R. Beach, Peter Kekenes-Huskey, Seth L. Robia

## Abstract

The ATP-dependent ion pump SERCA sequesters Ca^2+^ in the endoplasmic reticulum to establish a reservoir for cell signaling. Because of its central importance in physiology, this transporter is tightly controlled by physical interactions with tissue-specific regulatory micropeptides that tune SERCA function to match changing physiological conditions. In the heart, phospholamban (PLB) inhibits SERCA, while dwarf open reading frame (DWORF) stimulates SERCA. These competing interactions determine cardiac performance by modulating the amplitude of Ca^2+^ signals that drive the contraction/relaxation cycle. The distinct functions of these peptides may relate to their reciprocal preferences for SERCA binding. While SERCA binds PLB more avidly at low cytoplasmic Ca^2+^, it binds DWORF better at high Ca^2+^. In the present study, we determined that this opposing Ca^2+^ sensitivity is due to preferential binding of DWORF and PLB to different intermediate conformations that the pump samples during the Ca^2+^ transport cycle. The results suggest a mechanistic basis for inhibitory and stimulatory micropeptide function. In addition, fluorescence resonance energy transfer (FRET) measurements revealed dynamic shifts in SERCA-micropeptide binding equilibria during cellular Ca^2+^ elevations. The data suggest Ca^2+^-dependent dynamic exchange of inhibitory and stimulatory micropeptides from SERCA during the cardiac cycle. Together, these mechanisms provide beat-to-beat modulation of cardiac Ca^2+^ handling and contribute to the heart’s adaptation to the increased physiological demands of exercise.

**Significance** Interactions between SERCA and its regulatory micropeptides modulate cardiac performance. A previous study demonstrated that while the SERCA inhibitor, phospholamban, loses affinity during intracellular Ca^2+^ elevations, its competitor DWORF stimulates SERCA and binds better at elevated Ca^2+^. Here, we found this Ca^2+^-dependent difference in affinity is driven by preferential micropeptide binding to different SERCA conformations sampled during its enzymatic cycle. Tight binding to different conformations of the pump may underly the distinct mechanistic functions of these regulators. Lower affinity during alternating phases of the cardiac cycle may drive dynamic exchange of inhibitory and stimulatory micropeptides during contraction and relaxation. Our results reveal mechanisms that modulate cardiac Ca^2+^ handling on a beat-to-beat basis and help the heart adapt to exercise.

## INTRODUCTION

The type 2a sarco/endoplasmic reticulum Ca^2+^-ATPase (SERCA2a) is a P-type ion transporter responsible for sequestering cytoplasmic Ca^2+^ into the sarcoplasmic reticulum (SR) of cardiac muscle cells. The rate of Ca^2+^ transport by SERCA2a determines how quickly the heart muscle relaxes during the diastolic phase of the cardiac cycle as the ventricle is filling with blood. SERCA2a function also sets the amplitude of SR Ca^2+^ release, which determines the heart’s contractile strength during the systolic phase when blood is being ejected from the heart. Pathological decreases in SERCA expression, function, and regulation are associated with heart failure (6-8), focusing attention on SERCA as a possible therapeutic target (9, 10). So far, attempts to enhance transport function in patients by increasing expression of SERCA2a with gene delivery have been unsuccessful (11). Therefore, there is great interest in understanding physiological SERCA regulatory mechanisms to create a path toward rationally designed therapies that improve the function of endogenous SERCA.

The principal regulator of SERCA function in the heart is phospholamban (PLB), a single-pass transmembrane micropeptide (**Fig. 1**, blue). PLB physically interacts with SERCA and reduces its apparent affinity for Ca^2+^, decreasing Ca^2+^ uptake (12). This inhibition is relieved at high concentrations of cytoplasmic Ca^2+^ and after phosphorylation of PLB by cAMP-dependent protein kinase A (PKA) (13, 14), providing a mechanism to increase Ca^2+^ transport in response to exercise and other physiologic stress. The relief of SERCA inhibition was initially attributed to dissociation of the PLB-SERCA complex after PLB phosphorylation or Ca^2+^ binding to SERCA (15-19). However, subsequent spectroscopic studies showed that the PLB-SERCA interaction was constitutive (20, 21), with only modest changes in PLB-SERCA affinity in high Ca^2+^ and after PLB phosphorylation (22, 23). Thus, PLB may be considered a subunit of SERCA, and the mechanism for relief of inhibition must be due to some other structural rearrangement of the intact PLB-SERCA complex (24). We have previously shown that Ca^2+^-dependent shifts in SERCA conformational poise alter the apparent affinity of SERCA for PLB (4, 22, 25). We hypothesize PLB must bind more tightly to certain SERCA conformations, and this may be a key aspect of its regulatory function. However, the interaction energetics of PLB with major intermediate states in the Ca^2+^ transport cycle have not yet been measured. In the present study, we quantified PLB-SERCA interactions by FRET, controlling the conformational poise of SERCA while the enzyme was still in its intact cellular membrane environment.

**Figure 1.**
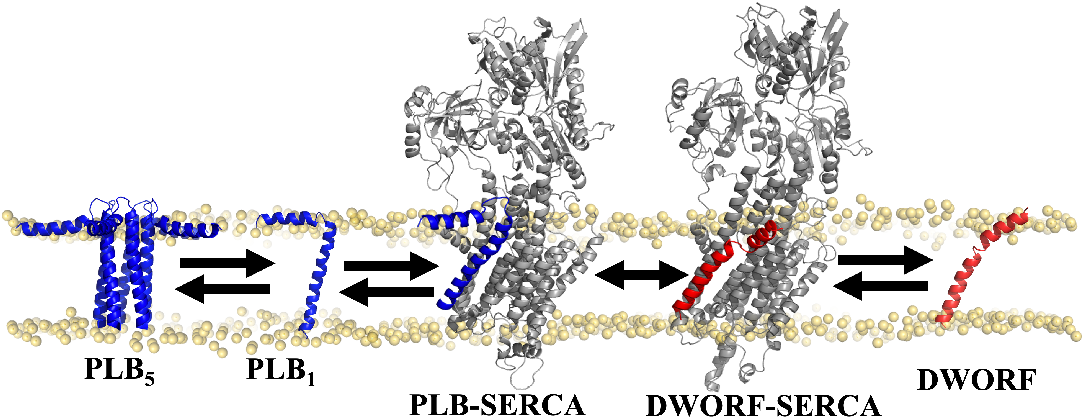
Regulatory interactions of SERCA (gray) with PLB (blue) and DWORF (red). Structures: PLB_5_ – PDB:2KYV (1); PLB_1_ – PDB:1FJP, (2); PLB-SERCA, DWORF-SERCA (3, 4); DWORF-7MPA, (5).

In addition to direct regulation by phosphorylation, PLB inhibitory potency may be modulated indirectly by oligomerization of PLB into pentamers (**Fig. 1**, “PLB_5_”). PLB pentamers represent an inactive, reserve pool (26) that serves as a buffer, reducing and stabilizing the concentration of the active, monomeric species. This buffering effect may be tuned by sympathetic stimulation, since oligomerization is further stabilized by PLB phosphorylation (23, 27). The physiological responsiveness of these dynamic binding equilibria must be governed by the rates of exchange of PLB protomers between SERCA- and pentamer-bound pools. We (28) and others (29) have provided evidence that the exchange of PLB monomers from pentamers occurs slowly relative to rapid exchange from the SERCA regulatory complex, but the underlying kinetics of these regulatory binding events have not been definitively measured. Therefore, the degree to which PLB monomers may dynamically redistribute among these pools in the cardiac SR remains unclear.

Here, we compared the binding dynamics of PLB with that of another SERCA-regulating membrane micropeptide expressed in the heart, dwarf open reading frame (DWORF) (**Fig. 1**, red) (5). In contrast to inhibitory PLB, DWORF has been proposed to increase Ca^2+^ transport in the heart by competing with PLB for its binding site on SERCA, relieving inhibition (30, 31), and it may also directly stimulate the pump (4, 32). We previously observed that DWORF has low oligomerization affinity, but DWORF and PLB have similar affinity for SERCA (33, 34). Interestingly, we recently determined that DWORF-SERCA binding shows an opposite Ca^2+^-dependence compared to PLB-SERCA. That is, DWORF affinity for SERCA increases with elevated Ca^2+^ (4). This implies that Ca^2+^ elevations favor DWORF in the competition with PLB for SERCA binding, but then PLB is favored as Ca^2+^ levels fall during diastole. The apparent dynamic competition of DWORF and PLB may be an important factor in future therapeutic strategies based on gene delivery of micropeptides (31, 35). We compared the energetics and kinetics of PLB and DWORF interactions with SERCA using a combination of enzymology and cell biology techniques. The results reveal mechanistic differences between inhibitory and stimulatory cardiac micropeptides and provide insight into how the heart may adapt to the shifting physiological demands of rest and exercise.

## RESULTS

### SERCA Affinity for PLB and DWORF is Dependent on Ca^2+^-pump conformation

We previously demonstrated that PLB-SERCA binding affinity is reduced by 30-40% in response to a sustained elevation of intracellular Ca^2+^, whereas the DWORF-SERCA interaction is slightly *more* stable with elevated [Ca^2+^] (4, 22). Since SERCA conformational poise changes between periods of elevated Ca^2+^ (during cardiac contraction) and low Ca^2+^ (during cardiac relaxation) (36), we reasoned that PLB and DWORF must preferentially bind to different intermediate conformations of the SERCA enzymatic cycle. The SERCA enzymatic cycle is represented in **Fig. 2A** with a simplified Post-Albers scheme (37). At basal Ca^2+^ (low nM), SERCA predominantly resides in the ATP bound state, E1-ATP (38) (**Fig. 2A**, blue box). During intracellular Ca^2+^ elevations, SERCA samples all physiological enzymatic states of the Ca^2+^ transport cycle, but the intermediate states are not all equally populated during systole. Rather, there is relative accumulation in the autophosphorylated E1P/E2P intermediates that precede rate-limiting steps in its enzymatic cycle (36, 39, 40). These states are outlined with red boxes in **Fig. 2A**.

**Figure 2.**
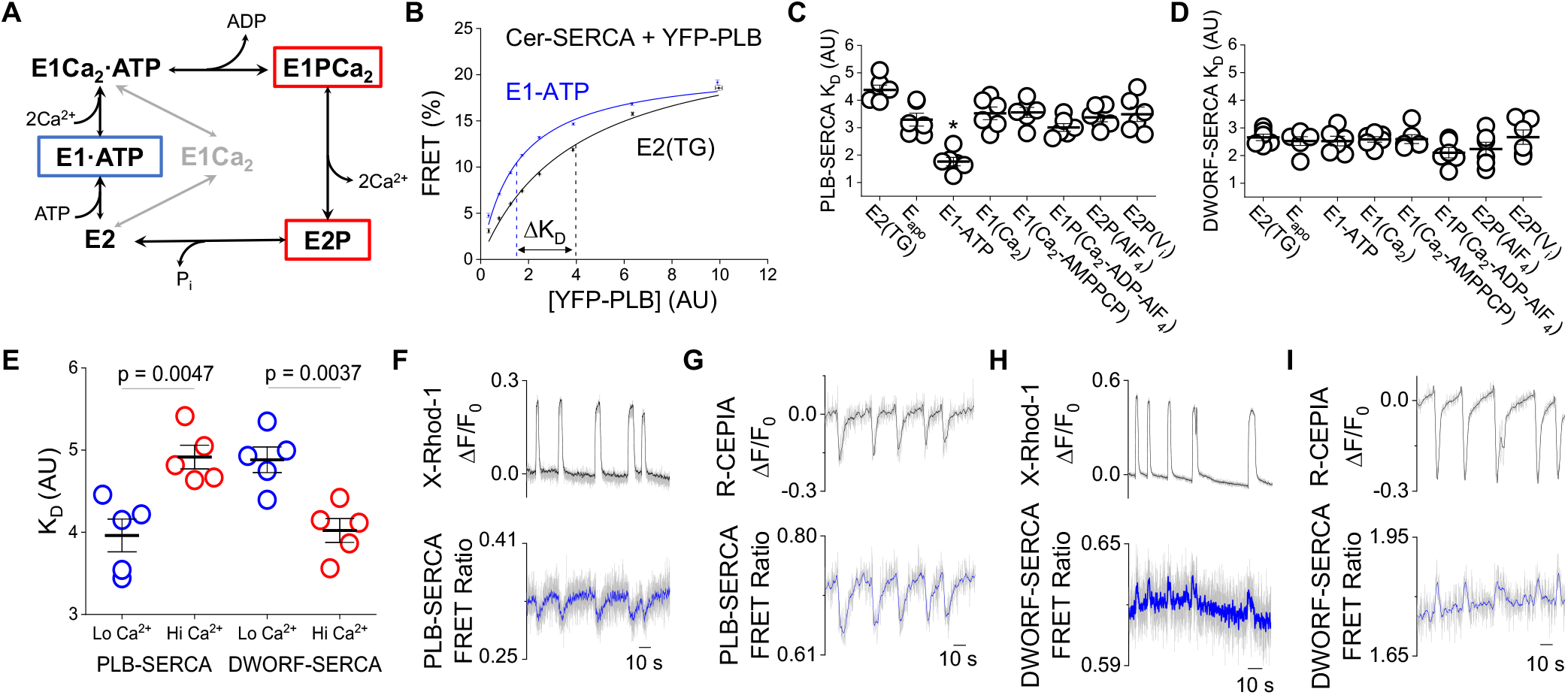
Dynamics of PLB and DWORF binding to SERCA during elevations in intracellular Ca^2+^. **A)** A simplified Post-Albers scheme of the SERCA enzymatic cycle, highlighting states that predominate at low (*blue*) and high (*red*) intracellular [Ca^2+^]. **B)** FRET-based binding curves displaying a shifted dissociation constant (K_D_) of PLB-SERCA binding between the ATP-(*blue*) and TG-bound (*black*) states of SERCA. **C**,**D)** Apparent K_D_s of PLB or DWORF binding to different SERCA enzymatic states of the catalytic cycle as in panel A with lines representing mean ± SE. Ligands used to stabilize each state are shown in parentheses. Differences in micropeptide K_D_s between SERCA states were analyzed by one-way ANOVA with Tukey’s *post-hoc* (*=p<0.05). **E)** Apparent K_D_s of PLB and DWORF for SERCA in ATP-containing solutions with low (*blue*) and high (*red*) concentrations of intracellular Ca^2+^ with lines representing mean ± SE. Differences in K_D_ evaluated by Student’s t-test. **F)** Confocal microscopy quantification of intracellular Ca^2+^ measured by X-rhod-1 fluorescence (*grey raw data, with black smoothed trendline*) with simultaneous measurement of changes in PLB-SERCA FRET (YFP/Cer ratio) (*grey raw data, with blue smoothed trendline*). **G)** Quantification of ER luminal Ca^2+^ measured by R-CEPIA1er fluorescence with simultaneous measurement of PLB-SERCA FRET (YFP/Cer ratio). **H)** Quantification of intracellular Ca^2+^ measured by X-rhod-1 fluorescence with simultaneous measurement of DWORF-SERCA FRET (YFP/Cer ratio) **I)** Quantification of ER luminal Ca^2+^ measured by R-CEPIA1er fluorescence with simultaneous measurement of DWORF-SERCA FRET (YFP/Cer ratio).

To determine the relative affinity of both micropeptides for these and other key SERCA enzymatic states, we transfected HEK-293 cells with Cerulean (Cer)-labeled SERCA2a (FRET donor) and yellow fluorescent protein (YFP)-labeled PLB (FRET acceptor) and quantified the interaction of these proteins using acceptor sensitization FRET microscopy. To control the conformational poise of SERCA, cells were permeabilized with 0.05 mg/mL saponin in bath solutions appropriate for stabilization of the transporter in various conformations (*see Methods*, **Fig. S1**). The affinity of the PLB-SERCA interaction was quantified by measuring FRET with automated fluorescence microscopy, as previously described (34, 41). FRET was low in cells with low fluorescence (low protein expression), increasing to a maximum in the brightest cells (high protein expression), yielding a FRET based “binding curve” (**Fig. 2B**). A hyperbolic fit to the data yielded the maximal FRET efficiency at high protein concentration or FRET_max_ (representing the intrinsic FRET of the bound PLB-SERCA complex) and the apparent dissociation constant (K_D_), the protein concentration that yields half-maximal FRET efficiency. The K_D_ value is inversely related to the affinity of the PLB-SERCA complex. **Figure 2B** shows the conditions that yielded the greatest difference in relative K_D_ for the PLB regulatory complex. PLB bound to the E1-ATP state of SERCA is obtained with a solution containing 3 mM ATP and low Ca^2+^, and represents the prevailing conformation in resting (diastolic) conditions. Addition of thapsigargin (TG) resulted in a significant right-shift of the binding curve (p = 8 × 10^−3^) (**Table S1**), indicating a decrease in PLB-SERCA affinity. Reduced affinity of PLB for TG-inhibited SERCA is consistent with previous observations from our lab (22) and others (19).

The summary of fitting of binding curves obtained from 6 independent transfections for each of 8 conditions is shown in **Fig. 2C-D** and **Table 1** (**Fig. S2** and **S3**). We observed especially avid binding of PLB to the E1-ATP state, which may indicate stabilization of that conformation of SERCA. This is the state that most SERCA pumps are in during the diastolic phase of the cardiac cycle when cytoplasmic Ca^2+^ is low, the muscle is relaxed, and the heart is filling with blood. The affinity of PLB for this state was not altered over a range of pH from 6-8 (**Fig S4**), indicating no preference for “E2” or “E1” transmembrane domain conformations stabilized in acidic or alkaline pH respectively (42, 43).

**Table 1.**
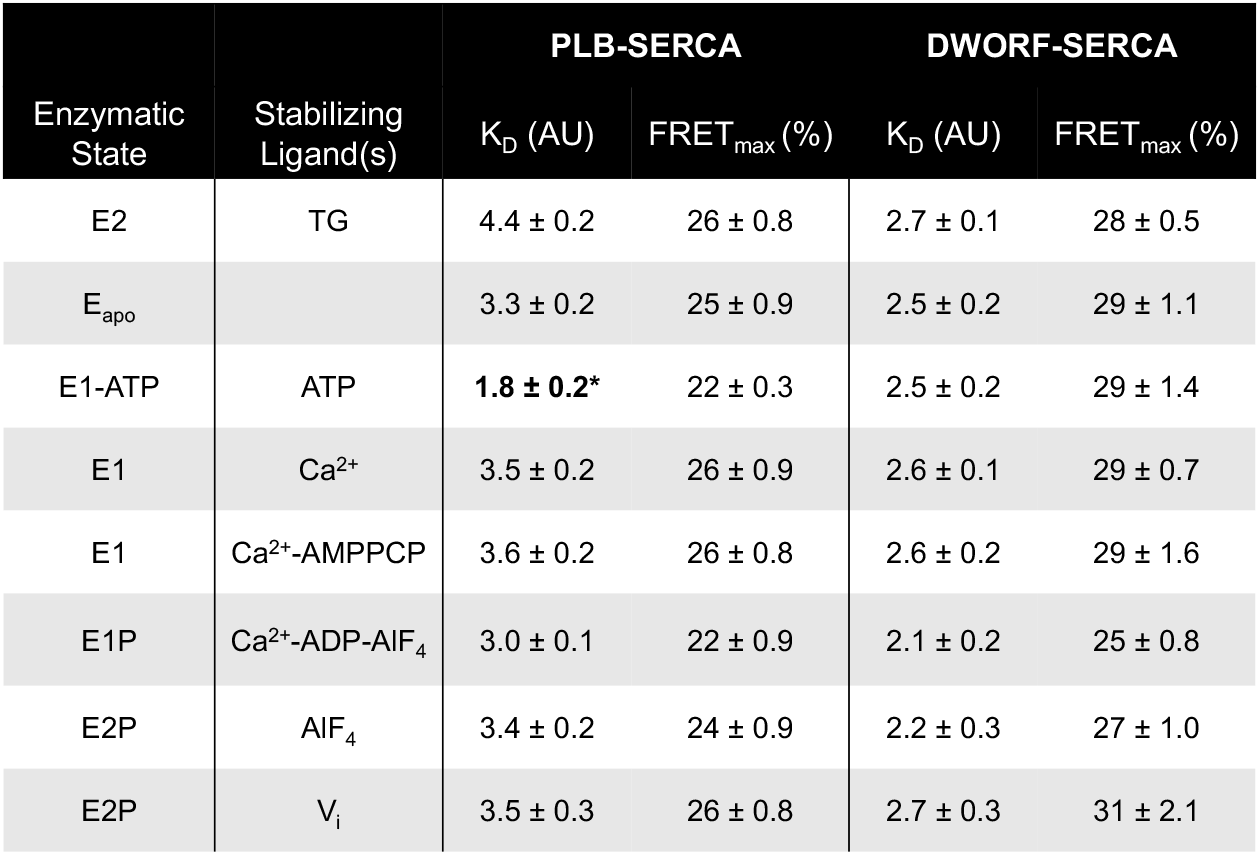
Acceptor-sensitization FRET binding curve parameters. Data are reported as mean ± SE. Differences were determined by one-way ANOVA with Dunn’s *post-hoc* (**\*** = p < 0.05, See Table S1 for complete statistical analysis).

In contrast to PLB, DWORF showed a much flatter affinity profile across SERCA enzymatic states (**Fig. 2D**). We noted slightly lower K_D_ values for E1PCa and E2P states. The population of SERCA pumps increasingly accumulates in these states during the systolic phase of the cardiac cycle when cytoplasmic Ca^2+^ is elevated and the heart is contracting to eject blood. DWORF may have a preference for those conformations, though the difference was not statistically significant by one-way ANOVA (**Table 1, Table S1**). We did resolve statistically significant differences in PLB-SERCA and DWORF-SERCA affinities in a simpler comparison of affinities in ATP-containing solutions: low Ca^2+^ “relaxing solution” vs. a high Ca^2+^ solution that induced enzymatic cycling of SERCA (*see Methods*). In these physiological buffers mimicking diastole vs. systole, respectively, we observed reciprocal Ca^2+^-dependence of binding affinity for PLB-SERCA vs. DWORF-SERCA (**Fig. 2E, S5**), consistent with our previous study (4).

Taken together, these data indicate that PLB and DWORF prefer to bind different SERCA conformations corresponding to different enzymatic states of the transport cycle. To determine how this differential Ca^2+^-dependent affinity may cause dynamic shifts in PLB and DWORF binding equilibria during intracellular Ca^2+^ elevations, we exploited a cardiomimetic model system described previously (44). HEK-293 cells expressing RyR and SERCA2a show spontaneous ER Ca^2+^ release events that give rise to large, prolonged increases in cytoplasmic Ca^2+^. Ca^2+^ transients were detected by confocal microscopy as an increase in X-rhod-1 cytoplasmic Ca^2+^ indicator fluorescence or a decrease in R-CEPIA1er ER Ca^2+^ indicator fluorescence (**Fig. 2F-I**, black). In some cells, these Ca^2+^ elevations coincided with small decreases in PLB-SERCA FRET, shown in **Fig. 2F** and **G** as grey data overlaid with a smoothed trendline (blue, *see Methods*) corresponding to a loss of PLB-SERCA affinity at elevated Ca^2+^. These were very small changes in FRET, as expected from the small differences in affinity quantified in equilibrium experiments (**Fig. 2C** and **E**). Indeed, we were previously unable to detect a PLB-SERCA FRET change over a single Ca^2+^ transient in electrically paced cardiac myocytes, only observing a modest decrease in average FRET over a period of repeated Ca^2+^ transients during rapid pacing (22). However, in HEK cells the Ca^2+^ elevations were more prolonged (increasing time for quantification), expression of endogenous SERCA/PLB was very low (reducing competing non-FRET interactions) and the cells are non-contractile (eliminating confounding cell motions). These advantages enabled detection of small shifts in the binding equilibria in the present study.

Next, we tested whether there was a change in DWORF-SERCA FRET in response to Ca^2+^ elevations. These FRET changes were even smaller than those observed for PLB-SERCA binding (**Fig. 2H**), consistent with DWORF’s relatively flat affinity profile across SERCA enzymatic states (**Fig. 2D**). The DWORF-SERCA FRET change was easier to appreciate in experiments where a low affinity Ca^2+^ indicator localized to the ER lumen (R-CEPIA1er) was used instead of a cytoplasmic Ca^2+^ indicator dye (**Fig. 2I**), perhaps because ER Ca^2+^ buffering by the indicator resulted in larger amplitude Ca^2+^ release events. Interestingly, the direction of the DWORF-SERCA FRET change was the opposite of that observed for PLB-SERCA; we observed modestly *increased* FRET during Ca^2+^ transients (**Fig. 2H** and **I**), consistent with enhanced DWORF-SERCA binding affinity at elevated cytoplasmic Ca^2+^ (**Fig. 2E**) (4).

### PLB Re-association with SERCA After Ca^2+^ Transients is Delayed by the PLB Pentamer

In examining the FRET changes accompanying Ca^2+^ elevations, we noted differences in the rates of recovery of FRET signals back to baseline after each Ca^2+^ transient, suggesting different rates of exchange of DWORF and PLB from SERCA. Specifically, during Ca^2+^ transient relaxation, DWORF appeared to dissociate rapidly from SERCA, as measured by the decrease in the DWORF-SERCA FRET ratio back to basal levels (**Fig. 2H** and **I**). However, PLB-SERCA re-association displayed a noticeable lag compared to the Ca^2+^ transient relaxation, with FRET continuing to increase back toward its resting maximum for several seconds after Ca^2+^ had already stabilized at a basal level (**Fig. 2F** and **G**). Exponential decay fitting of the recovering FRET signals and Ca^2+^ decay revealed characteristic time constants (τ) for each process (**Fig. 3, Table S2**). We found that PLB-SERCA re-association was significantly slower (τ = 4.9 ± 0.6 s) than the Ca^2+^ decay time (τ = 1.6 ± 0.2 s, p = 2 × 10^−8^) (**Fig. 3A, Table S3**). While the amplitude of the Ca^2+^-dependent DWORF-SERCA FRET change was very small (**Fig. 2H** and **3B**) due to its relatively flat affinity profile (**Fig. 2D**) we were able to quantify DWORF-SERCA unbinding (τ = 1.2 ± 0.4 s) and found it was similar to the Ca^2+^ transient relaxation time (**Fig. 3B**). **Fig. 3D** compares the rates of PLB-SERCA and DWORF-SERCA binding equilibrium shifts, expressed as the difference between the Ca^2+^ relaxation and the corresponding recovery of the FRET signal.

**Figure 3.**
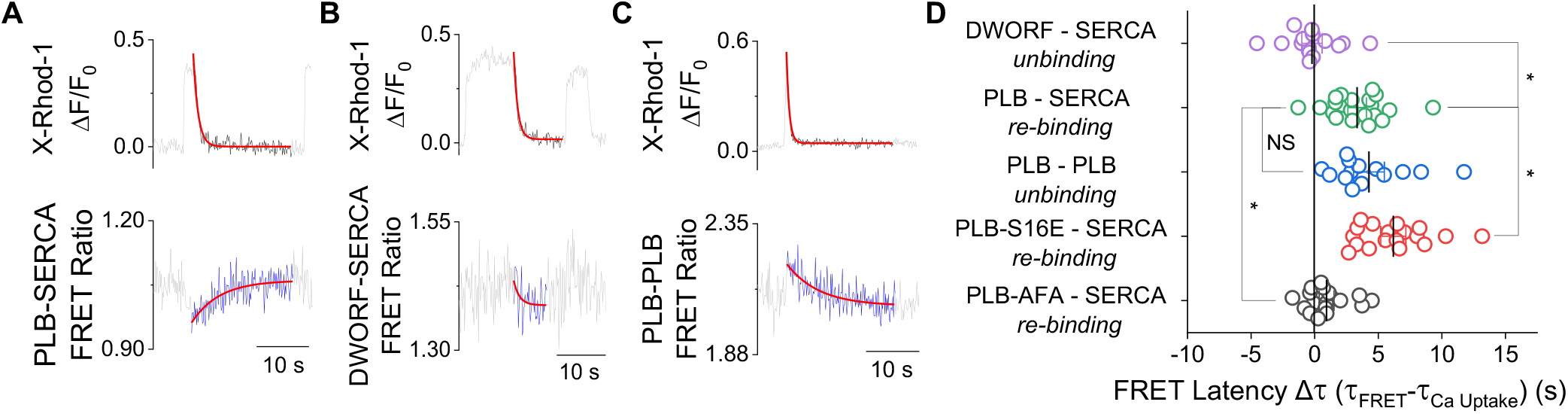
PLB re-association after Ca^2+^ transients is delayed by slow dissociation from the PLB pentamer. **A)** Kinetics of PLB-SERCA re-binding during Ca^2+^ uptake. **B)** Representative single exponential decay fit of the kinetics of DWORF-SERCA unbinding during Ca^2+^ uptake. **C)** Kinetics of PLB-PLB unbinding during Ca^2+^ uptake. **D)** The latency of FRET ratio changes compared to Ca^2+^ uptake with lines representing mean ± SE. Differences determined by one-way ANOVA with Dunn’s *post-hoc* test (* = p<0.05, see Table S4 for complete statistical analysis).

To understand the unexpectedly slow rate of PLB-SERCA re-binding, we considered whether the rate of PLB-SERCA recovery could be limited by slow dissociation of PLB protomers from the PLB pentamer (28, 29). To observe Ca^2+^-dependent changes in PLB oligomeric interactions, we transfected cells with Cer-PLB and YFP-PLB to measure intra-pentameric FRET. PLB-PLB FRET increased rapidly during Ca^2+^ elevations inversely to what was observed for PLB-SERCA FRET, indicating that PLB displaced from SERCA are redistributed into pentamers (**Fig. S6**). Notably, PLB-PLB FRET relaxation (pentamer unbinding) displayed a slow time course (τ = 5.2 ± 0.8 s) (**Fig. 3C**) that closely matched the slow PLB-SERCA reassociation (4.9 ± 0.6 s) (**Fig. 3D**). The data suggested PLB pentamer dissociation is indeed rate-limiting for rebinding of PLB to SERCA. Based on this observation, we predicted that PLB-SERCA re-binding rate would be changed by altering the stability of the PLB pentamer with mutagenesis. PLB pentamers are stabilized by PKA phosphorylation of serine 16 of PLB (27) during physiological stress, and this effect can be mimicked with S16E mutation (23). As predicted, stabilization of pentameric interactions with S16E mutation resulted in even slower PLB-SERCA reassociation (τ = 8.6 ± 0.7 s, p = 2 × 10^−6^) (**Fig. 3D** and **S6, Table S3**). Conversely, when the PLB pentamer was destabilized by alanine substitutions of 3 transmembrane cysteines (PLB-AFA) (8), PLB-SERCA re-association was significantly accelerated (τ = 2.5 ± 0.5 s, p = 8 × 10^−3^) (**Fig. 3D**, and **S6, Table S3**). The kinetics of SERCA binding/unbinding PLB and DWORF during Ca^2+^ uptake are summarized in Supplementary **Figure S7** and **Table S2**. Taken together, these results support the hypothesis that PLB oligomerization is the cause of the delayed re-association of the dynamic fraction of PLB monomers with SERCA after Ca^2+^ elevation. Moreover, the exaggerated delay of PLB reassociation with SERCA after S16E mutation suggests an adaptive physiological mechanism. Adrenaline-induced phosphorylation of PLB serine 16 may enhance the kinetic trapping (temporal buffering) of PLB monomers in pentamers *in-vivo* after adrenergic stimulation in conditions of physiologic stress.

We also investigated shifts in regulatory binding equilibria during the upstroke of the Ca^2+^ transient. This phase of the transient was too fast for adequate sampling using the previous confocal microscopy acquisition rate (7 images/sec), so we used a faster microscope to achieve an acquisition rate of 42 images/sec (*see Methods*). DWORF-SERCA binding occurred with a time course that was similar to that of Ca^2+^ release (τ = 0.22 ± 0.03 s and 0.21 ± 0.03 s, respectively) (**Fig. S8** and **Table S5**). However, PLB-SERCA unbinding was slower (τ = 0.74 ± 0.14 s), significantly lagging behind the Ca^2+^ upstroke (p = 4 × 10^−8^) (**Fig. S8** and **Table S5-7**). Likewise, we observed significantly slower PLB-PLB binding (τ = 0.59 ± 0.04 s) vs. Ca^2+^ release (p = 2 × 10^−4^) (**Fig S8** and **Table S5-7**) that was similar to PLB-SERCA unbinding time. The data suggest PLB-SERCA unbinding may be rate-limiting for subsequent PLB oligomerization during Ca^2+^ elevations.

### Modeling the Redistribution of PLB and DWORF Regulatory Complexes

The spontaneous Ca^2+^ elevations in HEK cells expressing RyR and SERCA2a are less frequent and longer in duration than cardiac Ca^2+^ transients, so we developed a computational model to integrate the measured rates of the dynamic interactions of PLB and DWORF with SERCA on the timescale of the cardiac cycle. The purpose of this model was to simulate the dynamic redistribution and exchange of regulatory species on a beat-to-beat basis and on the longer timescale between rest and exercise. The model describes the kinetics of these regulatory interactions with a set of numerically solved ordinary differential equations (see *Supplemental Methods*). A genetic algorithm was used to fit mean rate constants for the forward and reverse reactions in the model from experimentally measured FRET data as previously described (45). For simplicity, we considered the population of SERCA pumps to be distributed between two ensembles of pump enzymatic states, representing the “diastolic” condition (Ca-free) and the “systolic” condition (Ca-bound), as diagrammed in **Fig. 4A**. The relative population of these two ensemble states is dependent on the relative concentration of Ca^2+^, as quantified by confocal microscopy experiments. Monomeric PLB (PLB1) can bind either the Ca-free or Ca-bound ensemble of SERCA (22), but with higher affinity for the Ca-free population (**Fig. 2E**). The relative affinity of PLB for SERCA was constrained to be 2-fold higher for Ca-free vs. Ca-bound ensembles, as determined experimentally from FRET binding curves (**Fig. 2C**) (4). Monomeric PLB (PLB_1_) is also in equilibrium with the pentameric population (PLB_5_). The relative affinity of DWORF for SERCA was constrained to increase by 25% between Ca-free and Ca-bound ensembles, consistent with FRET measurements (**Fig. 2D**) (4). The computed kinetic parameters are provided in the Supplemental Data (**Table S8** and **S9**).

**Figure 4.**
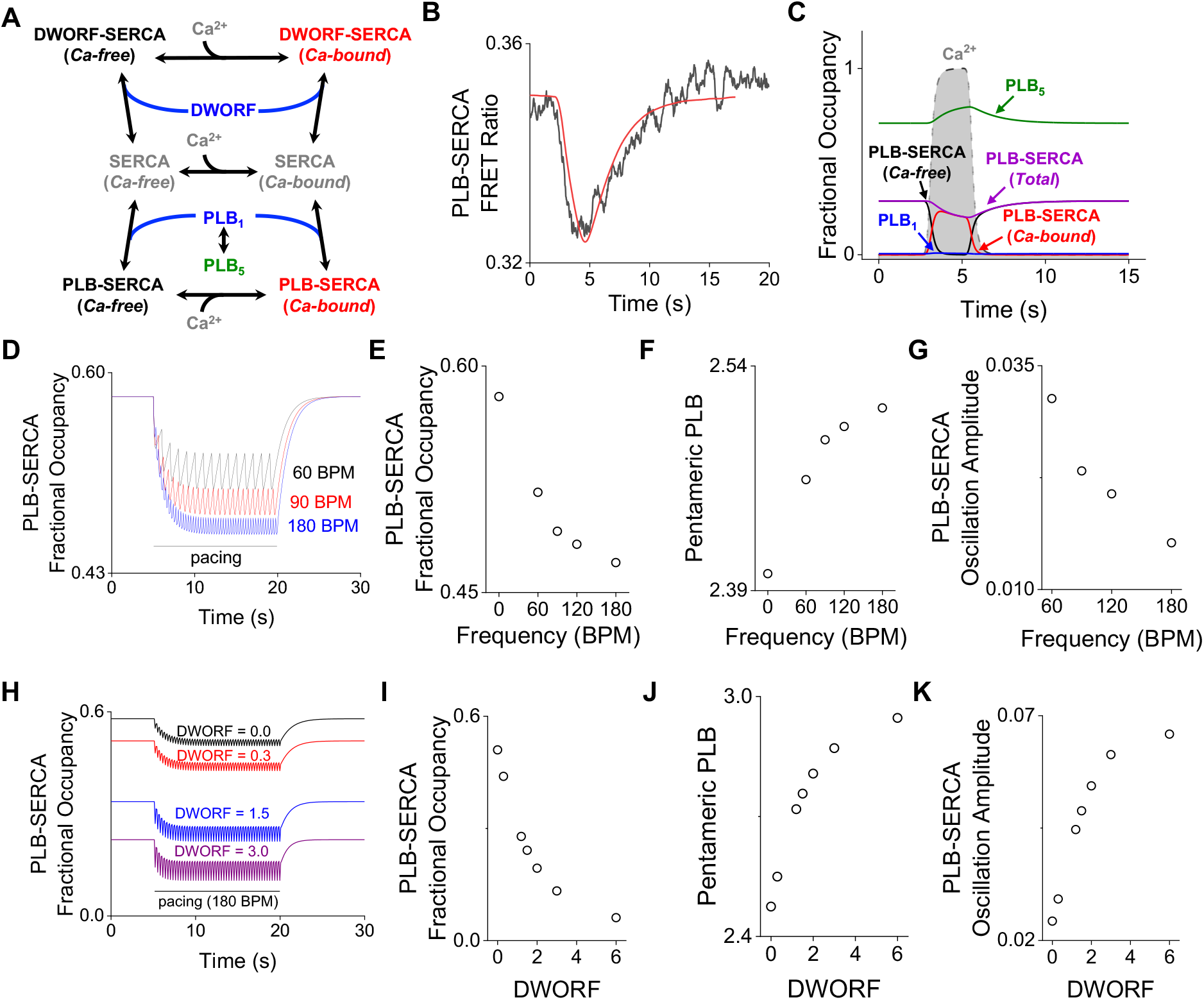
A computational model simulates the dynamics of PLB and DWORF interactions with SERCA. **A)** Simplified diagram of modeled regulatory interactions. **B)** A fit of the model to a representative FRET change measured by confocal microscopy. **C)** Simulation of changes in the populations of regulatory species during a Ca^2+^ elevation, where relative amounts of SERCA and PLB are equal. **D)** Simulation of the effect of cardiac pacing on PLB-SERCA at three heart rates (beats per min, BPM), where the ratio of SERCA:PLB was 1:3. **E)** Increasing pacing frequency modestly decreased PLB-SERCA binding. **F)** Increasing pacing frequency increased PLB oligomerization. **G)** PLB-SERCA oscillation amplitude decreased with faster pacing. **H)** Simulation of the effect of increasing DWORF expression relative to SERCA on PLB-SERCA binding. **I)** Increasing DWORF resulted in a decrease in the equilibrium level of PLB-SERCA. **J)** Increasing DWORF increased PLB oligomerization. **K)** Increasing DWORF relative to SERCA resulted in larger oscillations in PLB-SERCA binding. For H-K, the ratio of SERCA:PLB was 1:3 and pacing rate was 180 BPM.

### Dynamic Responses to Ca^2+^ Transients

The computational model provided good descriptions of data from physical experiments showing time-dependent changes in PLB-SERCA FRET and cytoplasmic Ca^2+^ (measured simultaneously). **Fig. 4B** provides a best-fit model prediction of the FRET compared to a representative experimental measurement. The model captured the rapid reduction of FRET with the rise of the Ca^2+^ transient. The decrease in PLB-SERCA is followed by a slower restoration of the FRET as Ca^2+^ returns to baseline. The similarity of the predicted and experimentally measured FRET data suggests that the model appropriately represents key aspects of the PLB-SERCA regulatory complex binding dynamics.

Using these fit parameter values, we simulated the time-dependent changes in PLB-SERCA and PLB-PLB binding dynamics that we observed in HEK cells. Importantly, we observed that a majority of the PLB-SERCA regulatory complexes remained intact (**Fig. 4C**, purple) during a Ca^2+^ elevation (**Fig. 4C**, grey shaded region), consistent with the “subunit model” (46). Thus, PLB can remain bound to SERCA when SERCA is bound to Ca^2+^; this Ca^2+^-bound PLB-SERCA fraction is highlighted in **Fig. 4C** (red). However, a fraction of the population of PLB-SERCA dissociated during a Ca^2+^ transient (**Fig. 4C**, purple). This fraction comprised ∼30% of PLB-SERCA complexes under these simulation conditions. This shift impacted the PLB monomer/pentamer equilibrium (**Fig. 4C**, blue/green). Expectedly, there was very little observed change in the small population of PLB monomers, as the dynamic fraction was quickly incorporated into pentamers. These reciprocal shifts are the cause of the transient decrease in SERCA-PLB FRET and increase in PLB-PLB FRET observed by confocal microscopy (**Fig. 3A** and **C**).

Next, we simulated the shorter, faster Ca^2+^ transients observed in the heart, pacing at a range of frequencies from 60 to 180 beats per minute, to investigate regulatory dynamics under conditions of rest, moderate exercise, and intense exercise. Increasing pacing frequency progressively decreased PLB-SERCA binding (**Fig. 4D, E**) and increased accumulation of PLB in pentamers (**Fig. 4F**). The systole-diastole difference in PLB-SERCA binding was small. This oscillation became even smaller with increasing pacing frequency (**Fig. 4D, G**), as the shifts in binding equilibria began to lag behind the rapid Ca^2+^ changes. These results suggest that the exchange dynamics of PLB may impact adaptive SERCA regulation between resting and exercising heart rates.

Introduction of DWORF-SERCA interactions to the model yielded the expected effect of competition of DWORF and PLB for SERCA binding (**Fig 4A, Table S9**). Increasing the amount of DWORF relative to PLB decreased PLB-SERCA binding (**Fig. 4H** and **I**) and increased PLB oligomerization (**Fig. 4J**). DWORF competes potently with PLB even at low stoichiometry because oligomerization of PLB reduces the effective concentration of the active monomeric species. Interestingly, the Ca^2+^-dependent increase in DWORF affinity for SERCA exaggerated the oscillations in PLB-SERCA binding during pacing (**Fig. 4K**). That is, during Ca^2+^ elevations DWORF (which binds SERCA better at high Ca^2+^) increasingly displaced PLB (which binds SERCA better at low Ca^2+^).

## DISCUSSION

### Differential Ca^2+^-dependence of micropeptides for SERCA

The principal finding of the present study is that the opposite Ca^2+^-dependence of DWORF and PLB binding to SERCA (4) (**Fig. 2E**) is due to differential affinity of these competing micropeptides for particular enzymatic states in the transport cycle (**Fig. 2C** and **2D**). The FRET-based binding assay showed that PLB has the highest affinity for the E1-ATP state, which is the predominant state at resting Ca^2+^ when the cardiac muscle is relaxed and the ventricle is filling with blood (cardiac diastole). We consider micropeptide-SERCA binding affinity to be indicative of the relative energetics of the PLB-SERCA complex, much as melting temperature may be quantified as a proxy for the stability of a protein-protein complex (47). Stabilization of the E1-ATP structure is expected to favor the population of that state, slowing the subsequent Ca^2+^-binding step. This is expected to reduce the apparent Ca^2+^ affinity of the pump and pump turnover, the principal inhibitory effects of PLB on SERCA (12). In contrast, DWORF shows a much flatter SERCA-binding profile, with more avid binding to E1P and E2P states. Much of the SERCA population accumulates in these states when cytoplasmic Ca^2+^ is elevated and the heart is contracting to eject blood (cardiac systole). This accumulation occurs during rapid Ca^2+^ transport as a result of rate-limiting steps in the enzymatic cycle (39, 40) that create “traffic jams”. There are two mechanistic implications of the observed tight binding of DWORF to these states. If DWORF stabilizes these states (or adjacent intermediates) it may reduce an activation energy barrier that limits the kinetics of these enzymatic transitions. This could accelerate pump turnover, an effect that would be in harmony with recent studies that suggest a direct *stimulatory* effect of DWORF on SERCA ATPase activity and Ca^2+^ uptake rate (4, 5, 32). Secondly, DWORF may compete more effectively for SERCA-binding during cytoplasmic Ca^2+^ elevations, increasingly displacing inhibitory PLB and exaggerating the dynamic fraction that unbinds at high Ca^2+^. Dynamic competition of these regulatory peptides is noteworthy, since the discovery of DWORF raised the question of why SERCA regulation should involve both an inhibitor (PLB) and an inhibitor of the inhibitor (DWORF). This seems cumbersome if the mechanistic goal is simply more or less Ca^2+^ transport activity. We reasoned that the interplay of these opposing regulators must be necessary for responsiveness to changing conditions. This could include reacting to transient Ca^2+^ changes during the cardiac cycle or on a longer timescale, responding to adrenergic signaling and changes in pacing frequency during exercise. The present data provide insight into Ca^2+^ transport regulation on both timescales:

### Functional implications of beat-to-beat micropeptide exchange

The opposite Ca^2+^-dependence of SERCA binding to DWORF and PLB (4) (**Fig. 2C-E**) is noteworthy in the context of cardiac physiology. Importantly, PLB-SERCA complexes do not dissociate *en masse* during systole (**Fig. 4D**); we have previously reported that elevated Ca^2+^ causes only small shifts in regulatory complex binding equilibria (4, 22). Still, cardiac Ca^2+^ transients simultaneously reduce PLB binding affinity and favor DWORF binding, driving a “dynamic fraction” of the population of these micropeptides to exchange from SERCA during each Ca^2+^ transient. The functional impact of this exchange is expected to be an exaggeration of the intrinsic response of SERCA to changing Ca^2+^. That is, during diastole the already low SERCA activity is further inhibited by an increase in the fraction of pumps that bind PLB (**Fig. 2F** and **G**). During systole, high SERCA activity is further boosted by enhanced DWORF binding (**Fig. 2H** and **I**), which increasingly displaces PLB and directly stimulates SERCA maximal activity (4, 32). Together, low Ca^2+^ inhibition and high Ca^2+^ stimulation should enhance the apparent cooperativity of the Ca^2+^ response, which would benefit cardiac function by conserving ATP consumption until Ca^2+^ transport is most efficient and most necessary.

A second anticipated effect of dynamic competition of the micropeptides is a phase shift of the SERCA activation/inhibition cycle relative to cytoplasmic Ca^2+^ oscillations. Specifically, while PLB-SERCA unbinding is relatively fast, we did observe a modest delay between Ca^2+^ release and unbinding of the dynamic fraction of PLB (**Fig. S8**) suggesting this inhibitory interaction persists briefly during the initial rise in Ca^2+^. Thus, slow dissociation of the dynamic fraction of PLB transiently protects the micropeptide binding site from DWORF. Consequently, in the presence of a slowly dissociating dynamic fraction of PLB, full SERCA stimulation by DWORF also lags in time compared to the Ca^2+^ upstroke. Together, a persistent fraction of PLB and a lagging DWORF transiently decrease SERCA activity compared to the level of transport that would be supported by that level of Ca^2+^ under equilibrium conditions. The functional consequence of this lag is to prevent Ca^2+^ uptake from opposing Ca^2+^ release early in systole. This is expected to increase the amplitude and rate of rise of Ca^2+^ transient and minimize futile cycling of the pump at peak Ca^2+^, thereby conserving ATP resources. Conversely, during the onset of diastole, kinetic trapping of PLB in slowly dissociating PLB pentamers (**Fig. 3C**) results in a significant lag between the Ca^2+^ transient decay and rebinding of PLB to SERCA (**Fig. 3A**). Slow rebinding of PLB to SERCA is expected to prolong elevated SERCA activity, making the Ca^2+^ decline steeper and yielding a lower concentration of cytoplasmic Ca^2+^ during diastole.

### Micropeptide dynamics during rest and exercise

The observations presented here also have implications for the cardiac stress response. During the transition from rest to exercise, cardiac Ca^2+^ handling accelerates in response to more frequent Ca^2+^ elevations (increased heart rate) and adrenergic stimulation (PLB phosphorylation). The slow kinetics of PLB exchange from pentamers means that faster pacing should integrate more pentameric PLB at the expense of the population of PLB-SERCA (**Fig. 4D**). Thus, one may speculate that this mechanism contributes to a positive force-frequency relationship (treppe) (48). Moreover, during exercise, oligomerization is secondarily enhanced by PLB phosphorylation by PKA, which further slows the unbinding of PLB from oligomers during diastole (**Fig. 3D**). This would be reinforced by CaMKII activation by elevated Ca^2+^ during fast pacing. Interestingly, computational modeling showed that competitive binding of SERCA by DWORF decreases both the basal level of PLB-SERCA binding as previously proposed (30) and also increases the amplitude of the oscillations in the population of the PLB-SERCA complex (**Fig. 4I-K**). The dose-dependence of these effects revealed in the computational model may provide insight into DWORF-based gene therapy approaches.

### Summary and Future Directions

Dynamic competition of regulatory micropeptides on seconds and minutes timescales represents an important aspect of the responsiveness of SERCA regulation to Ca^2+^ transients and PLB phosphorylation, respectively. The data also clarify the role of the PLB pentamer as a phosphorylation-tunable kinetic trap that limits the rate of SERCA-rebinding by a dynamic fraction of PLB. Future studies will elucidate the functional predictions derived from the present results, specifically: DWORF enhances the E1P-E2P transitions in the SERCA enzymatic cycle; DWORF increases SERCA cooperativity in the presence of PLB; DWORF increases the small fraction of PLB that binds/unbinds SERCA with each Ca^2+^ elevation, exaggerating the SERCA inhibition/stimulation cycle; DWORF causes a phase shift in the SERCA inhibition/stimulation cycle relative to the cardiac Ca^2+^ oscillations. Overall, the results suggest the importance of dynamic competition of PLB and DWORF for cardiac function and may provide insight into the interplay of other tissue-specific micropeptide regulators of ion transporters more generally.

## ACKNOWLEDGEMENTS

The authors thank Howard S. Young, Gianluigi Veglia, J. Michael Autry, Daniel Koch, Catherine Makarewich, and L. Michel Espinoza-Fonseca for helpful discussions. The authors thank Elisa Bovo and Aleksey Zima for technical assistance. The authors thank L. Michel Espinoza-Fonseca for providing the minimized structures in Fig. 1. This investigation was supported by the National Institutes of Health (NIH): Maximizing Investigators’ Research Awards (MIRA) R35GM124977 from the National Institute of General Medical Sciences (NIGMS) to PK-H; MIRA R35GM138183 from NIGMS and K22HL133150 from the National Heart, Lung, and Blood Institute (NHLBI) to JB; R01HL092321 and R01HL143816 from the NHLBI to SLR.

## METHODS

### Plasmid Constructs

For all plasmid constructs, we used pEGFP-C1 as the expression vector in mammalian cells. All micropeptides (PLB, DWORF, PLB-AFA, and PLB-S16A) and SERCA constructs consisted of mCerulean3, EYFP, or TagRFP fused via a 5 amino acid linker to the N-terminus of the micropeptide or SERCA (28, 33, 49).

### Cell culture and transfection

AAV 293 cells were cultured in DMEM cell culture medium supplemented with 10% fetal bovine serum (FBS) (ThermoScientific, Waltham, MA). Following culture, the cells were transiently transfected using either MBS mammalian transfection kit (Agilent Technologies, Stratagene, La Jolla, CA) or Lipofectamine 3000 transfection kit (Invitrogen, Carlsbad, CA) as per instructions provided with the respective kits. 24 hours post-transfection the cells were trypsinized (ThermoScientific, Waltham, MA) and replated onto poly-D-lysine coated glass bottom chambers and allowed to settle down for one hour before imaging. We have previously estimated that this approach achieves endoplasmic reticulum protein expression levels on the order of 2000:1 or 4000:1 lipids:protein ratio (50), vs. 750:1 lipid:protein ratio quantified for native sarcoplasmic reticulum (46, 51). Thus, native protein concentrations are 3-5 fold more concentrated and we do not consider this heterologous model to represent “overexpression” of SR proteins.

### FRET Acceptor Sensitization in Permeabilized HEK-293 cells

Acceptor sensitization FRET was quantified as previously described (22). Briefly, AAV 293 cells were transiently transfected with Cer-donor and YFP-acceptor labeled FRET binding partners in a 1:5 molar plasmid ratio, and FRET acceptor sensitization was measured by automated fluorescence microscopy before and after permeabilization with 0.005% w/v saponin. For each condition, 2 sets of 72 images (∼500 total cells per condition) were collected from six independent experiments with a 20X 0.75 N.A. objective with 50 ms exposure for each channel: Cer, YFP, and FRET (Cer excitation/ YFP emission). Cells that expressed Cer, had an area of 136-679 µm^2^, and were at least 40% circular were automatically scored for Cer, YFP, and FRET fluorescence intensity with a rolling background subtraction using a plugin in ImageJ. FRET efficiency was calculated according to *E* = *G* /(*G* + 2.782 X *F*_*Cer*_), where *G* = *F*_*FRET*_ − *a* X *F*_*YFP*_ − *d* X *F*_*Cer*_, where *F*_*FRET*_, *F*_*YFP*_, and *F*_*Cer*_ are the fluorescence intensities from FRET, YFP, and Cer images, respectively, and *G* represents FRET intensity corrected for the bleedthrough of the channels. The parameters *a* and *d* are bleedthrough constants calculated as *a* = *F*_*FRET*_ /*F*_*YFP*_ for a control sample transfected only with YFP-SERCA and *d* = *F*_*FRET*_ /*F*_*Cer*_ for a control sample transfected only with Cer-SERCA. For our experimental setup, *a* and *d* were 0.185 and 0.405 respectively. FRET efficiency for each scored cell was plotted as a function of expressed protein concentration, as determined from the fluorescence intensity of the yellow fluorescent protein channel (34, 41). FRET was low in cells with low fluorescence (low protein expression), increasing to a maximum in the brightest cells (high protein expression), yielding a FRET based “binding curve”. The data were fit to a hyperbolic function of the form [FRET = FRET_max_ [YFP]/(K_D_+[YFP]), where FRET_max_ is the maximal FRET efficiency at high protein concentration (representing the intrinsic FRET of the bound complex), [YFP] is inferred from YFP fluorescence emission in each cell, and K_D_ is the apparent dissociation constant (the protein concentration that yields half-maximal FRET efficiency).

To control the conformational poise of SERCA, cells were permeabilized with 0.05 mg/mL saponin in bath solutions appropriate for stabilization of the transporter in various conformations. Solutions for ligand-stabilization of enzymatic intermediate states were prepared by addition of corresponding substrates to a calcium-free base solution, which includes 100 mM KCl, 5 mM MgCl_2_, 2mM EGTA, and 10 mM imidazole, pH 7.0. The base solution was used to characterize SERCA in a ligand-free state, E_apo_. The following ligands were used to prepare specific solutions corresponding to SERCA biochemical state: 100 µM thapsigargin for E2(TG); 3 mM ATP for E1-ATP; 2.1 mM CaCl_2_ for E1(Ca_2_) with free[Ca^2+^]_i_ = 100 µM (52); 2.1 mM CaCl_2_ and 500 µM nonhydrolyzable ATP analog (AMPPCP) for E1(Ca_2_-AMPPCP); 2.1 mM CaCl_2_, 500 µM ADP, 50 µM AlCl_3_, and 3 mM KF for E1(Ca_2_–ADP–AlF_4-_); 0.1 mM orthovanadate for E2(Vi); and 50 µM AlCl_3_ and 3 mM KF for E2(AlF_4_^-^). Concentrations of AMP-PCP nucleotide analog and ADP nucleotide were reduced from previously published conditions (3 mM) used to stabilize SERCA in microsomal fractions (36) in order to prevent altering fluorescent protein emission intensity. The data for each set of binding partners were analyzed for differences in K_D_ between buffer conditions using a one-way ANOVA with Tukey’s *post-hoc* test (significance = p<0.05).

For experiments measuring PLB binding preference for SERCA enzymatic states E2-ATP and E1-ATP, which are in equilibrium under low Ca^2+^, high ATP conditions (e.g. cardiac diastole), a buffer containing 100 mM KCl, 5 mM MgCl_2_, 2mM EGTA, 10 mM imidazole, and 3 mM ATP was prepared. The pH was then adjusted to the following concentrations: pH = 6.0, 6.5, 7.0, 7.5, and 8.0. For these experiments, a TagRFP-PLB acceptor construct (pK_a_=3.8) was used in place of the YFP-PLB acceptor because YFP fluorescence is sensitive to pH changes in this range (pK_a_ = 6.9) (53).

For experiments with low and elevated intracellular [Ca^2+^], solutions were prepared containing potassium aspartate 120 mM, KCl 15 mM, KH_2_PO_4_ 5 mM, MgCl_2_ 0.75 mM, dextran 2%, ATP 5 mM, HEPES 20 mM, and EGTA 2 mM, pH 7.2. The elevated [Ca^2+^] buffer was prepared with CaCl_2_ 1.7 mM for a free [Ca^2+^]_i_ = 2 µM (52). The data for each set of binding partners were analyzed for differences in K_D_ between high and low Ca^2+^ conditions using a Student’s t-test (significance = p<0.05).

### Confocal fluorescence microscopy to measure intermolecular FRET and HEK-293 cytoplasmic Ca^2+^

HEK-293 cells exhibiting spontaneous Ca^2+^ oscillations were generated by transient transfection with GFP-tagged RyR2 and either Cer or unlabeled SERCA2a, and co-transfected with SERCA, PLB, PLB-AFA, PLB-S16E, or DWORF FRET pairs tagged with Cer and YFP fluorophores. Transfected cells were cultured for 24 h and seeded into poly-D-lysine-coated glass-bottom chamber slides in Dulbecco’s modified Eagle’s medium plus 10% fetal bovine serum. 24 h after seeding, cell culture medium was changed with PBS (+Ca^2+^/+Mg^2+^), and experiments were conducted with a Leica SP5 laser scanning confocal microscope equipped with a 63X water objective. To observe transient changes in cytoplasmic calcium, cells were incubated with 10 µM X-Rhod-1/AM (X-Rhod) for 20 min in PBS (+Ca^2+^/+Mg^2+^). X-Rhod was excited with the 543-nm line of a He-Ne laser, and emitted fluorescence was measured at wavelength 580 nm. FRET pair fluorophores Cer and YFP were excited with the 430 and 514 nm lines of an argon laser, respectively, and emitted fluorescence was measured at wavelengths 485 ± 15 and 537 ± 15 nm, respectively. Images were acquired in line-scan with averaging of 4 every 134 ms for ∼2 min.

A select set of concurrent experiments were conducted with a Zeiss LSM 880 Airyscan confocal microscope using a 40X oil immersion objective. X-Rhod was excited with the 594-nm line of a HeNe laser, and emitted fluorescence was measured at wavelength 580 nm. FRET pair fluorophores Cer and YFP were excited with the 458 nm line of an argon laser, and emitted fluorescence was measured at wavelengths 485 ± 15 and 537 ± 15 nm, respectively. Images were acquired in line-scan every 24 ms for ∼2 min.

FRET ratio was determined by dividing the acceptor fluorescence by the donor fluorescence and plotted as a function of time with X-Rhod to indicate concurrent changes in [Ca^2+^]. FRET ratio data was smoothed using a Savitzky-Golay binomial filter with a 4.08 s averaging window. Changes in FRET ratio and X-Rhod fluorescence associated with Ca^2+^ uptake were fit to the single-exponential decay function, *y* = *A*1^−*x*/*τ*^ + *y*_0_ in Origin, to estimate the time constant or τ of the change, where *A*1 is the amplitude of change and *y*_0_ is the initial detected fluorescence. Changes in FRET ratio and X-Rhod fluorescence associated with Ca^2+^ release were fit using the single-exponential decay function, 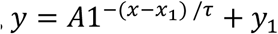, where *x*_*1*_ is the time in seconds of Ca^2+^ release, and *y*_*1*_ is the baseline fluorescence prior to *x*_*1*_, defined by the linear function *y* = *A*1 + *y*_2_. Differences in the characteristic time constants (τ) for each process were analyzed using a one-way ANOVA with Dunn’s *post-hoc* test (significance = p<0.05).

### Kinetic Modeling

We implemented systems of ordinary differential equations according to the schematic provided in Fig. 4A (54). This kinetic diagram describes the populations of the SERCA states under diastolic and systolic conditions. Model parameters were informed or constrained by experimental observations where appropriate, such as the PLB dissociation rates reported in this study. Mean rates were determined by averaging over several transients from multiple cells. Kinetic parameters for PLB-SERCA binding dynamics were fit to 15 independent FRET transients, while kinetic parameters for DWORF-SERCA interactions were fit to 9 independent FRET measurements. All other parameters were assigned initial values that were subject to fitting against time-varying FRET data. The relative affinity of PLB for SERCA (here described by the rate fraction of k_on_/k_off_) was constrained to be 2-fold higher for Ca-free vs. Ca-bound ensembles, as determined experimentally from FRET binding curves. Likewise, the relative affinity of DWORF for SERCA (k_on_/k_off_) was constrained to increase by 25% between Ca-free and Ca-bound ensembles, consistent with FRET measurements. The ODE system was numerically solved using the scipy (v1.5.0) ODEINT function, using the experimentally measured intracellular Ca^2+^ transient as an input to the model. The resulting numerically estimated PLB-SERCA population was compared against the experimentally reported FRET data by assuming that the PLB-SERCA population was proportional to the FRET efficiency. Mean-squared error between the predicted PLB-SERCA population and the FRET data was computed to assess the fit. Fitting was optimized using a genetic algorithm we developed previously (45) that reduces the mean-squared error by randomizing model parameters and selecting those that reduce error. Forward simulations of the SERCA/PLB/DWORF equilibria for the cardiac pacing experiments used simulated Ca^2+^ transients with amplitudes and frequencies in place of those measured in HEK cells for fitting purposes. All code written in support of this publication are publicly available at https://github.com/huskeypm/pkh-lab-analyses.

