## Supplementary Materials for "Inhibitory and Stimulatory Micropeptides Preferentially Bind to Different Conformations of the Cardiac Calcium Pump"

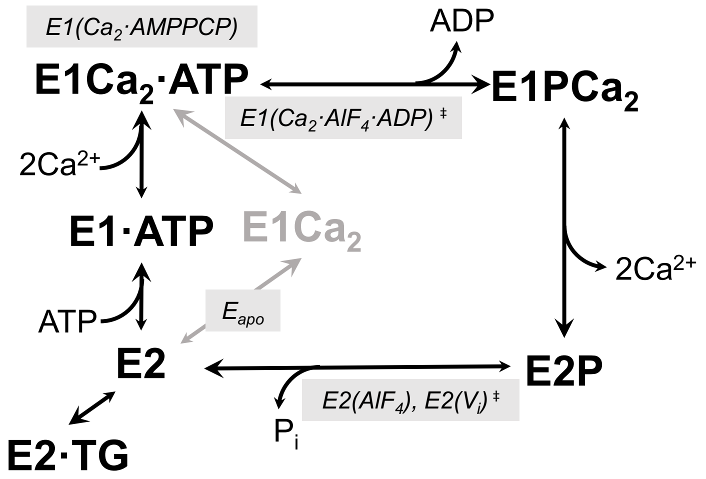


**Supplemental Figure S1.** Post-Albers cycle with biochemically stabilized states of SERCA. Ligand-free SERCA (pH 7) is in the $E_{apo}$ state – a state in pH-dependent dynamic equilibrium between E2 and E1 states where substrate binding sites face the ER lumen or the cytosol respectively (1). The SERCA inhibitor, thapsigargin (TG), stabilizes a dead-end E2 conformation (1IWO) (2). Saturating concentrations of ATP or $Ca^{2+}$ stabilize E1 conformations of the pump (3, 4). The structural analog of $E1Ca_2$-ATP where both substrates are bound is stabilized with $Ca^{2+}$ and a non-hydrolysable ATP-analog, AMPPCP (PDB: 1T5S) (5). The ADP-bound transition state following ATP-hydrolysis is stabilized with saturating concentrations of $Ca^{2+}$, ADP, and the phosphate analog, $AlF_4$ (PDB: 2ZBD) (6). A low $Ca^{2+}$ buffer containing either $AlF_4$ or the SERCA inhibitor orthovanadate ($V_i$) stabilize a transitional analog of the dephosphorylating E2P state (PDB: 3N5K, 5A3Q) (7, 8). ‡ indicates transition states.

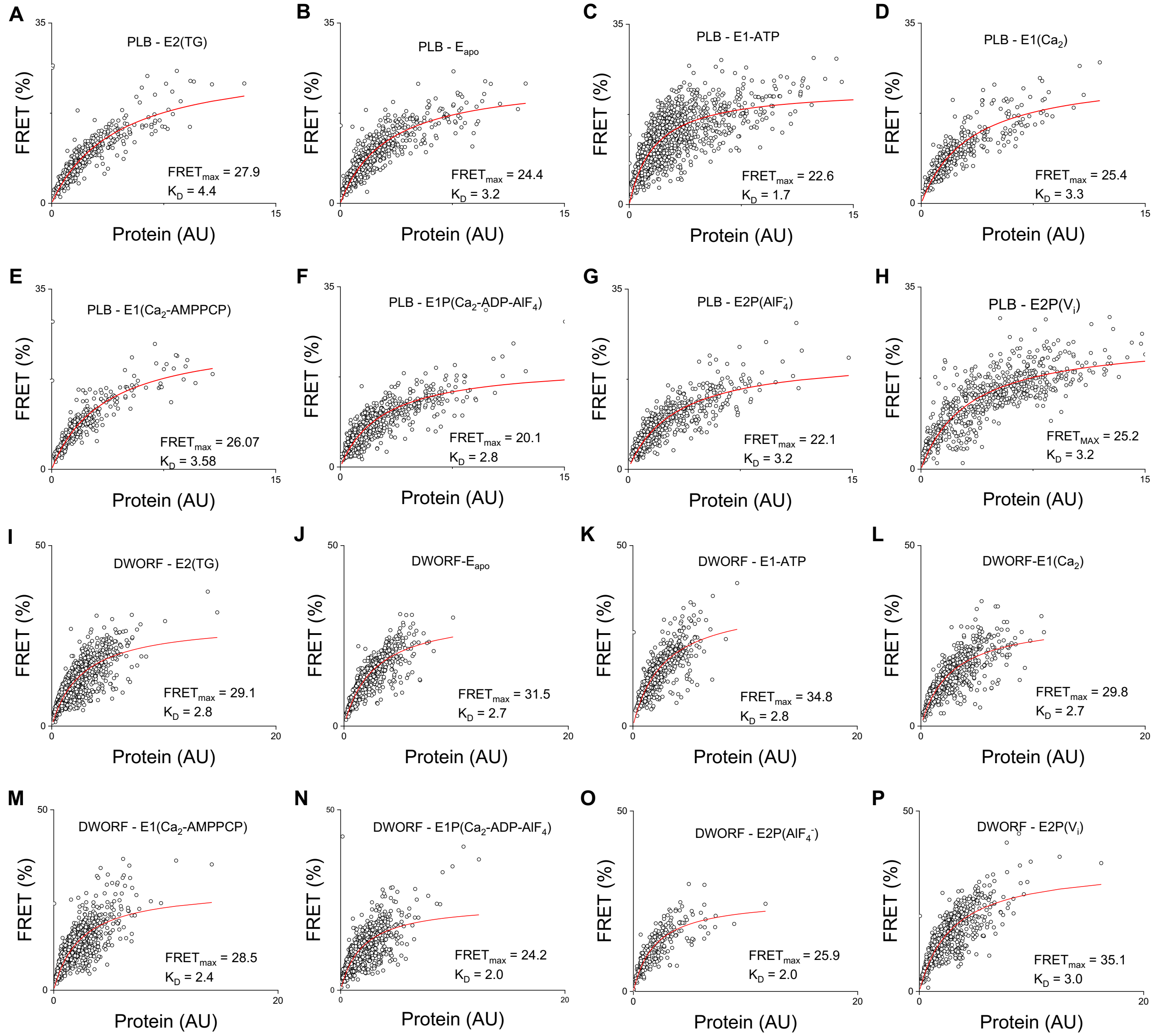


**Supplementary Figure S2.** Representative FRET-based binding curves for PLB and DWORF interactions with ligand-stabilized enzymatic states of SERCA. **A-H)** Representative binding curves based on FRET measurements between Cer-SERCA and YFP-PLB FRET pairs. **I-P)** Representative binding curves based on FRET measurements between Cer-SERCA and YFP-DWORF FRET pairs.

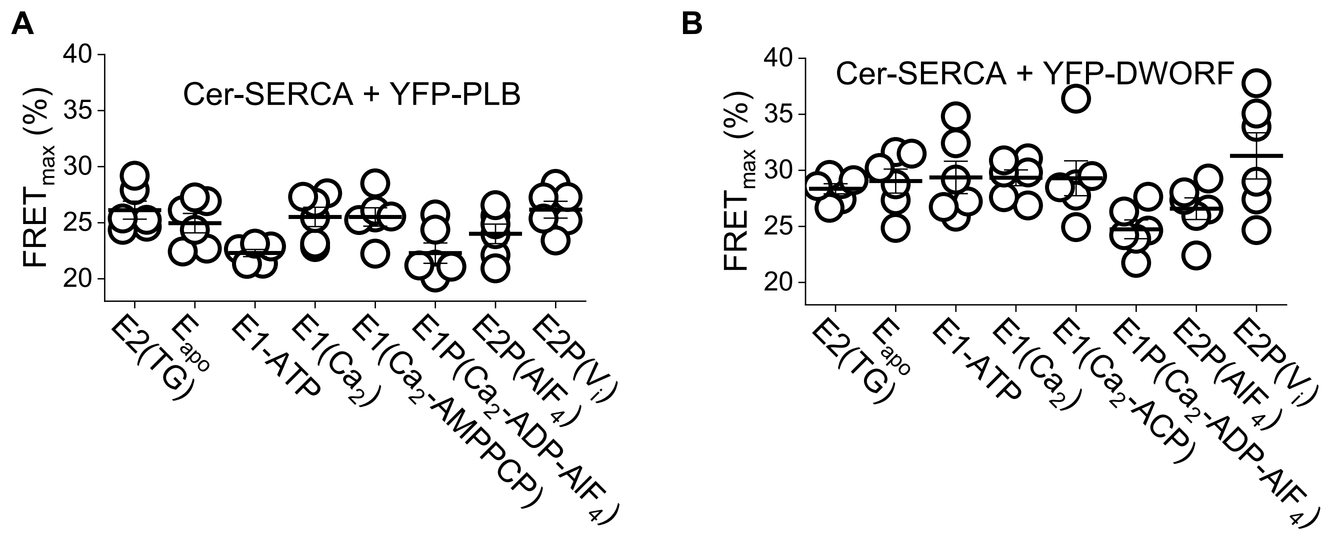


**Supplementary Figure S3.** $FRET_{max}$ values for PLB and DWORF regulatory complexes with ligand-stabilized enzymatic states of SERCA with lines representing mean ± SE. Data are derived from hyperbolic fits of FRET data plotted over a range of YFP-tagged micropeptide protein concentration (See representative curves in Supplementary Fig. S1).

| PLB-SERCA Conformation $K_D$ 1-way ANOVA with Tukey's post-hoc | | | | | | | |
|---|---|---|---|---|---|---|---|
| | E2(TG) | $E_{apo}$ | **E1-ATP** | E1($Ca_2$) | E1($Ca_2$-AMPPCP) | E1P($Ca_2$-ADP-$AlF_4$) | E2P($AlF_4$) |
| E2P($V_i$) | $5.14 \times 10^{-2}$ | 1 | **$7.07 \times 10^{-6}$*** | 1 | 1 | 0.67 | 1 |
| E2P($AlF_4$) | **$1.86 \times 10^{-2}$*** | 1 | **$2.54 \times 10^{-5}$*** | 1 | 1 | 0.89 | |
| E1P($Ca_2$-ADP-$AlF_4$) | **$3.93 \times 10^{-4}$*** | 0.97 | **$1.57 \times 10^{-3}$*** | 0.59 | 0.52 | | |
| E1($Ca_2$-ACP) | $9.13 \times 10^{-2}$ | 0.98 | **$3.33 \times 10^{-6}$*** | 1 | | | |
| E1($Ca_2$) | $6.90 \times 10^{-2}$ | 0.99 | **$4.76 \times 10^{-6}$*** | | | | |
| **E1-ATP** | **$1.96 \times 10^{-8}$*** | **$6.63 \times 10^{-5}$*** | | | | | |
| $E_{apo}$ | **$8.14 \times 10^{-3}$*** | | | | | | |

| DWORF-SERCA Conformation $K_D$ 1-way ANOVA with Tukey's post-hoc | | | | | | | |
|---|---|---|---|---|---|---|---|
| | E2(TG) | $E_{apo}$ | E1-ATP | E1($Ca_2$) | E1($Ca_2$-AMPPCP) | E1P($Ca_2$-ADP-$AlF_4$) | E2P($AlF_4$) |
| E2P($V_i$) | 1 | 1 | 1 | 1 | 1 | 0.39 | 0.71 |
| E2P($AlF_4$) | 0.73 | 0.95 | 0.95 | 0.88 | 0.86 | 1 | |
| E1P($Ca_2$-ADP-$AlF_4$) | 0.41 | 0.73 | 0.73 | 0.59 | 0.56 | | |
| E1($Ca_2$-ACP) | 1 | 1 | 1 | 1 | | | |
| E1($Ca_2$) | 1 | 1 | 1 | | | | |
| E1-ATP | 1 | 1 | | | | | |
| $E_{apo}$ | 1 | | | | | | |

**Supplementary Table S1.** P values comparing differential PLB (above) and DWORF (below) dissociation constants for SERCA conformation states. Data were analyzed by 1-way ANOVA with Tukey's *post-hoc* test ($p<0.05$ = *).

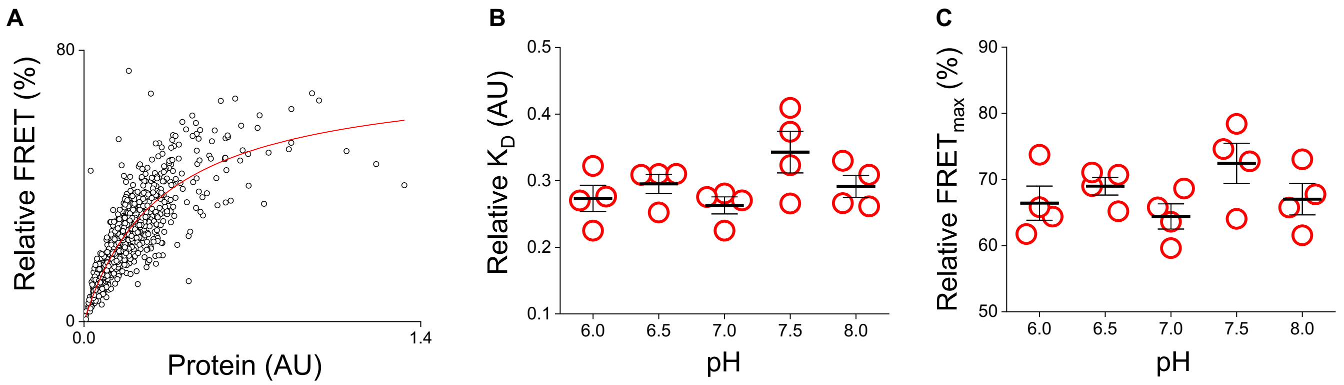


**Supplementary Figure S4.** PLB binding preference for E2-ATP and E1-ATP substates of the ATP bound state of SERCA was assessed by measuring Cer-SERCA – TagRFP-PLB FRET in a buffer containing ATP (3 mM) at a range of pH from 6-8 (*See Methods*). Increasingly acidic conditions protonate SERCA substrate binding sites, stabilizing E2 conformations, whereas E1 is favored at alkaline pH (1). **A)** Representative FRET-based binding curve for TagRFP-PLB interacting with ATP-bound Cer-SERCA at pH 7.0. **B)** Relative $K_D$ values of the PLB-SERCA regulatory complex measured at a range of pH from 6-8, with lines representing mean ± SE. We saw no change in PLB-SERCA $K_D$ in this range, indicating no discernable preference of PLB for E2-ATP or E1-ATP states. **C)** Relative $FRET_{max}$ values of the PLB-SERCA regulatory complex measured at a range of pH from 6-8, with lines representing mean ± SE.

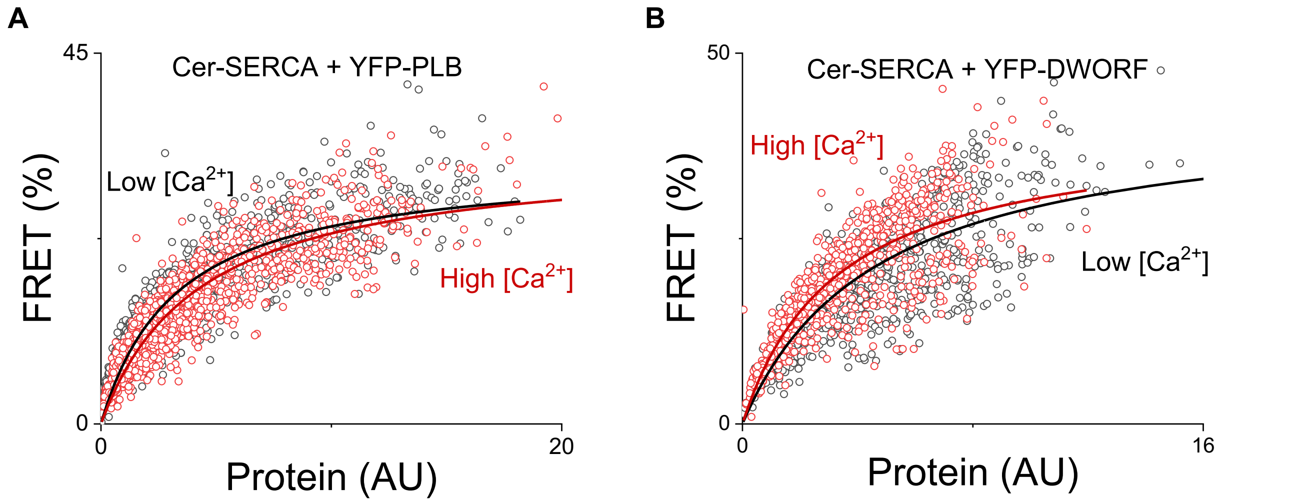


**Supplementary Figure S5.** Representative PLB-SERCA and DWORF-SERCA FRET-based binding curves in high and low [$Ca^{2+}$] solutions simulating phyioslogical diastolic and systolic conditions in permeabilized HEK-293 cells (*See Methods*).

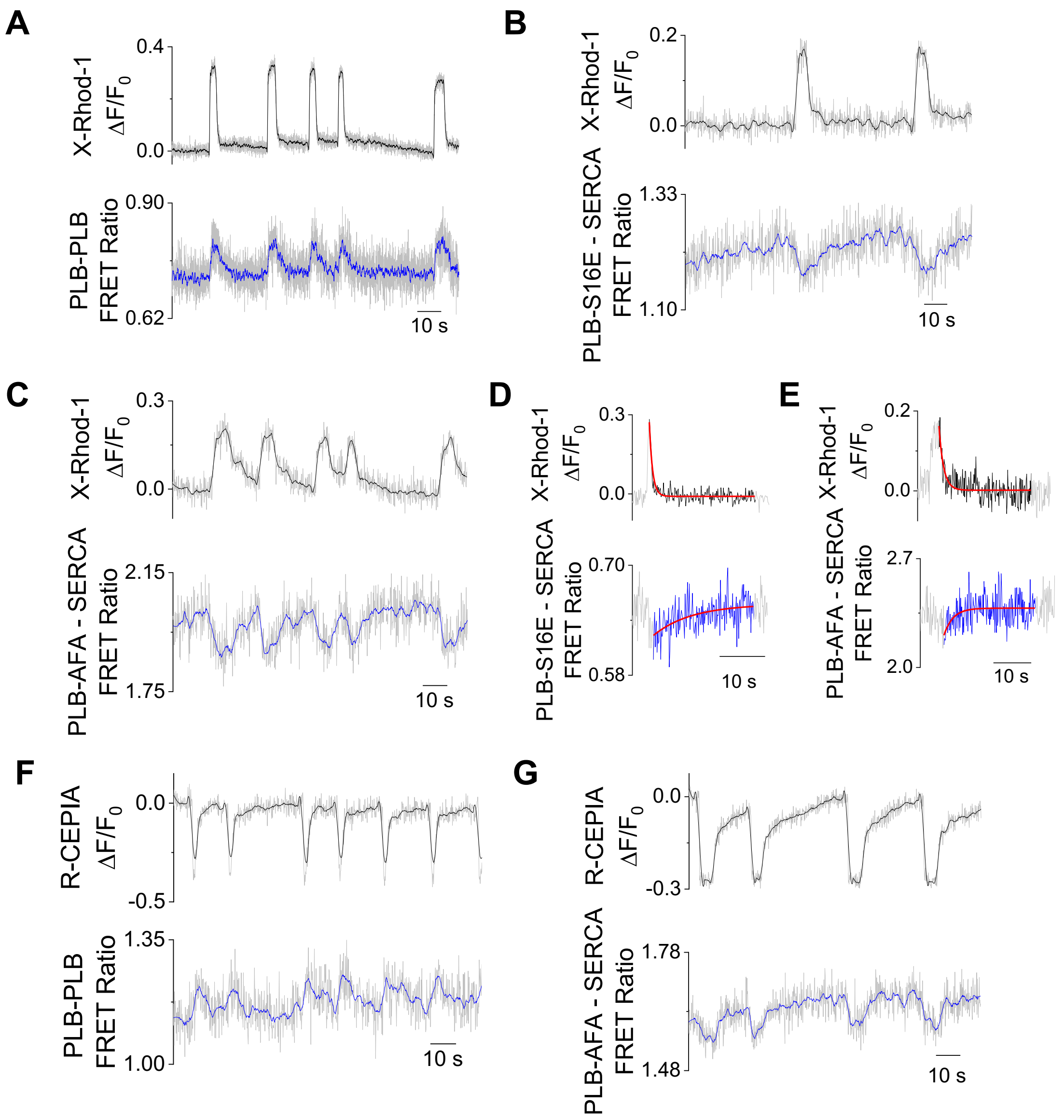


**Supplementary Figure S6.** Representative confocal microscopy data for PLB - PLB, PLB-S16E – SERCA, and PLB-AFA – SERCA FRET ratio (*grey + blue smoothed trendline*) changes measured in response to spontaneous intracellular $Ca^{2+}$ elevations measured by X-rhod-1 or RCEPIA1er fluorescence (*grey + black smoothed trendline*). **A)** Representative data trace of PLB-PLB FRET ratio transiently increasing in response to $Ca^{2+}$ elevations. **B)** Representative data trace of PLB-S16E – SERCA FRET ratio transiently decreasing in response to $Ca^{2+}$ elevations. **C)** Representative data trace of PLB-AFA – SERCA FRET ratio decreasing in response to $Ca^{2+}$ elevations. **D)** Representative single exponential decay fit of PLB-S16E – SERCA re-binding associated with $Ca^{2+}$ uptake. **E)** Representative single exponential decay fit of PLB-AFA – SERCA re-binding associated with $Ca^{2+}$ uptake. **F)** Representative data trace of PLB-PLB FRET ratio transiently increasing during ER $Ca^{2+}$ release. **G)** Representative data trace of PLB-AFA – SERCA FRET ratio transiently decreasing during ER $Ca^{2+}$ release.

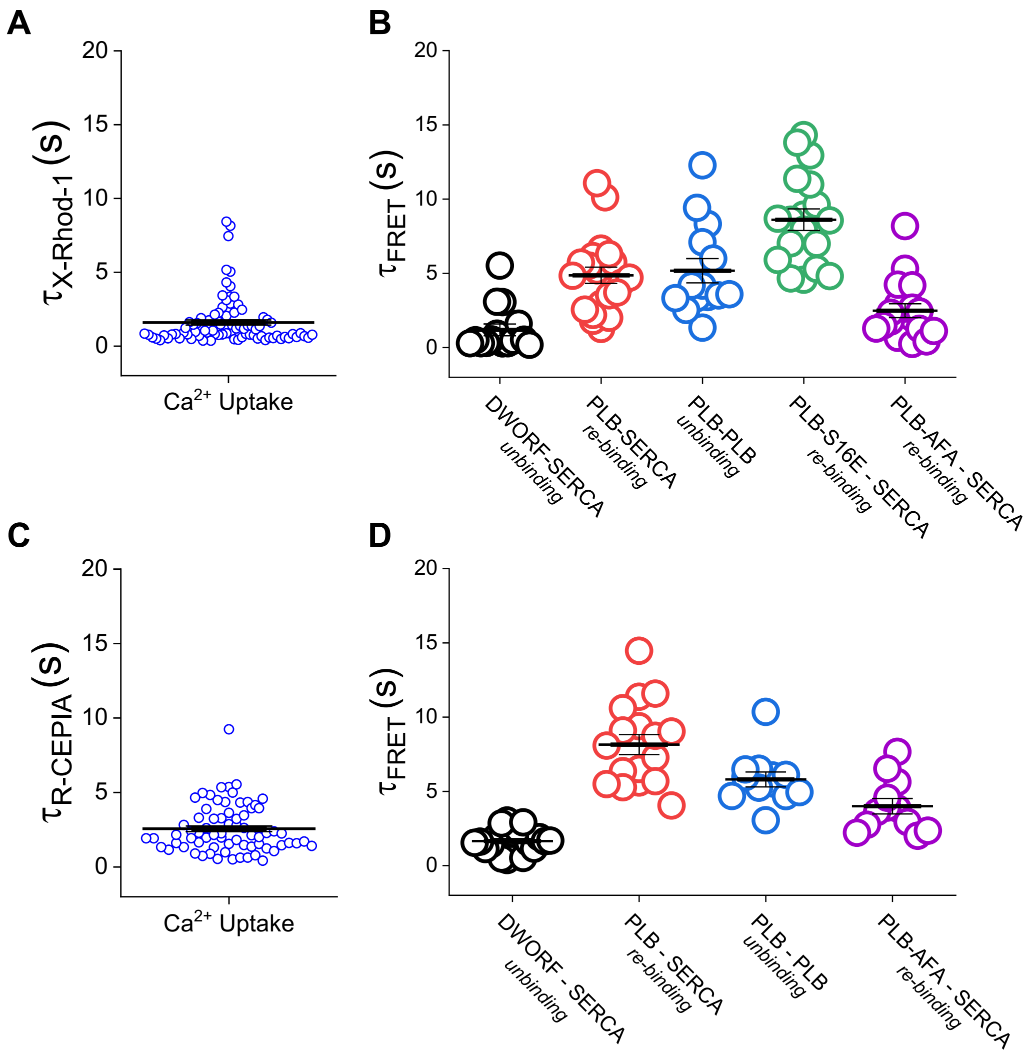


**Supplementary Figure S7.** Time constants (τ) quantified for regulatory binding processes associated with $Ca^{2+}$ uptake with lines representing mean ± SE. **A)** Time constant values for $Ca^{2+}$ uptake derived from single exponential decay fits of decreases in X-rhod-1 fluorescence. **B)** Raw time constant values for SERCA and micropeptide binding processes derived from single exponential decay fits of FRET ratio changes associated with $Ca^{2+}$ uptake measured by X-Rhod-1 fluorescence (data are shown in Fig. 3D as the FRET latency quantified by the difference between the τ of FRET ratio changes and the τ their corresponding $Ca^{2+}$ uptake event). See Table S3 for complete statistical analysis. **C)** Time constant values for $Ca^{2+}$ uptake derived from single exponential decay fits of increases in R-CEPIA1er fluorescence. **D)** Raw time constant values for SERCA and micropeptide binding processes derived from single exponential decay fits of FRET ratio changes associated with $Ca^{2+}$ uptake measured by R-CEPIA1er fluorescence.

| Apparent τ (Mean ± SE) | |
|---|---|
| Process | τ (s) |
| DWORF – SERCA (*unbinding)* | 1.2 ± 0.4 |
| PLB – SERCA (*re-binding)* | 4.9 ± 0.6 |
| PLB – PLB (*unbinding)* | 5.2 ± 0.8 |
| PLB-S16E – SERCA *(re-binding)* | 8.6 ± 0.7 |
| PLB-AFA – SERCA *(re-binding)* | 2.5 ± 0.5 |
| $Ca^{2+}$ Uptake | 1.6 ± 0.2 |

**Supplementary Table S2.** Time constants (τ) of binding processes associated with $Ca^{2+}$ uptake measured by X-Rhod-1 fluorescence. Time constant values are reported as mean ± SE.

| SERCA/Micropeptide binding/unbinding τ 1-way ANOVA with Dunn's post-hoc | DWORF-SERCA (*unbinding*) | PLB-SERCA (*re-binding*) | PLB-PLB (*unbinding*) | PLB-S16E - SERCA (*re-binding*) | PLB-AFA – SERCA (*re-binding*) |
|---|---|---|---|---|---|
| $Ca^{2+}$ Uptake | 1 | **$2.38 \times 10^{-8*}$** | **$2.81 \times 10^{-7*}$** | **$<1 \times 10^{-10*}$** | 0.81 |
| PLB-AFA - SERCA (*re-binding*) | 0.72 | **$7.95 \times 10^{-3*}$** | **$6.30 \times 10^{-3*}$** | **$<1 \times 10^{-10*}$** | |
| PLB-S16E - SERCA (*re-binding*) | **$<1 \times 10^{-10*}$** | **$1.64 \times 10^{-6*}$** | **$1.30 \times 10^{-4*}$** | | |
| PLB - PLB (*unbinding*) | **$1.35 \times 10^{-5*}$** | 1 | | | |
| PLB - SERCA (*re-binding*) | **$9.79 \times 10^{-6*}$** | | | | |

**Supplementary Table S3.** P values comparing differences in time constants (τ) for SERCA-micropeptide binding/unbinding processes analyzed by 1-way ANOVA with Dunn's *post-hoc* test. See Fig. S7A and B and Table S2 for summary data.

| FRET Latency Δτ ($\tau_{FRET} - \tau_{Ca\ Uptake}$) 1-way ANOVA with Dunn's post-hoc test | DWORF-SERCA (*unbinding*) | PLB-SERCA (*re-binding*) | PLB-PLB (*unbinding*) | PLB-S16E - SERCA (*re-binding*) |
|---|---|---|---|---|
| PLB-AFA - SERCA (*re-binding*) | 0.85 | **$2.24 \times 10^{-2*}$** | **$1.73 \times 10^{-3*}$** | **$3.48 \times 10^{-8*}$** |
| PLB-S16E - SERCA (*re-binding*) | **$2.96 \times 10^{-10*}$** | **$3.59 \times 10^{-3*}$** | 0.23 | |
| PLB-PLB (*unbinding*) | **$2.45 \times 10^{-5*}$** | 0.95 | | |
| PLB-SERCA (*re-binding*) | **$3.03 \times 10^{-4*}$** | | | |

**Supplementary Table S4.** P values comparing FRET latency Δτ ($\tau_{FRET} - \tau_{Ca\ Uptake}$) differences between groups analyzed by 1-way ANOVA with Dunn's *post-hoc* test. See Fig. 3D for summary data.

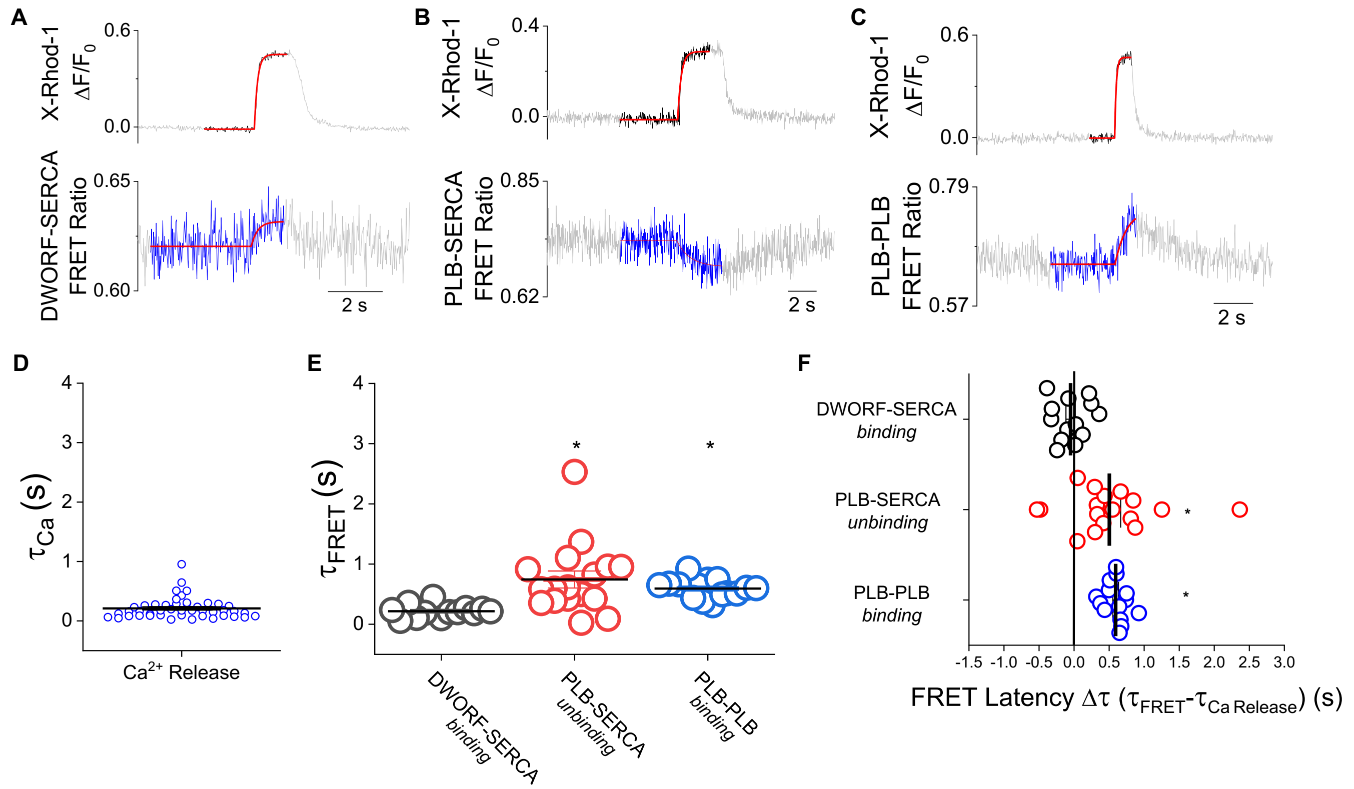


**Supplementary Figure S8.** PLB pentamer formation during $Ca^{2+}$ elevations is modestly delayed by the time course of PLB-SERCA unbinding. **A)** Representative single exponential decay fit of DWORF-SERCA binding during $Ca^{2+}$ release. **B)** Representative single exponential decay fit of PLB-SERCA unbinding during $Ca^{2+}$ release. **C)** Representative single exponential decay fit of PLB-PLB binding during $Ca^{2+}$ release. **D)** Time constant values for $Ca^{2+}$ release derived from single exponential decay fits of increases in X-rhod-1 fluorescence. **E)** Time constant values for SERCA and micropeptide binding processes derived from single exponential decay fits of FRET ratio changes associated with $Ca^{2+}$ release with lines representing mean ± SE. Differences determined by one-way ANOVA with Dunn's *post-hoc* test (* = $p<0.05$, see Table S6 for complete statistical analysis). **F)** The latency of FRET ratio changes lagging behind the kinetics of $Ca^{2+}$ release with lines representing mean ± SE. Differences determined by one-way ANOVA with Dunn's *post-hoc* test (* = $p<0.05$, see Table S7 for complete statistical analysis).

| Apparent τ (Mean ± SE) | |
|---|---|
| Process | τ (s) |
| DWORF – SERCA (*binding*) | 0.22 ± 0.03 |
| PLB – SERCA (*unbinding*) | 0.74 ± 0.14 |
| PLB – PLB (*binding*) | 0.59 ± 0.04 |
| $Ca^{2+}$ Release | 0.21 ± 0.03 |

**Supplementary Table S5.** Time constants (τ) quantified for regulatory binding processes associated with $Ca^{2+}$ release. Time constant values are reported as mean ± SE.

| SERCA/micropeptide binding/unbinding processes analyzed by 1-way ANOVA with Dunn's post-hoc | | | |
|---|---|---|---|
| | DWORF-SERCA (*binding*) | PLB-SERCA (*unbinding*) | PLB-PLB (*binding*) |
| $Ca^{2+}$ Release | 1 | **3.84 x $10^{-8}$*** | **1.74 x $10^{-4}$*** |
| PLB-PLB (*binding*) | **5.79 x $10^{-3}$*** | 0.62 | |
| PLB-SERCA (*unbinding*) | **2.45 x $10^{-5}$*** | | |

**Supplementary Table S6.** P values comparing differences in time constants (τ) for SERCA/micropeptide binding/unbinding processes analyzed by 1-way ANOVA with Dunn's *post-hoc* test. See Fig. S8D and E and Table S5 for summary data.

| FRET Latency Δτ ($\tau_{FRET} - \tau_{Ca\ Uptake}$) 1-way ANOVA with Dunn's post-hoc test | | |
|---|---|---|
| | PLB-SERCA (*unbinding*) | PLB-PLB (*binding*) |
| DWORF-SERCA (*binding*) | **4.06 x $10^{-3}$*** | **1.08 x $10^{-3}$*** |
| PLB-PLB (*binding*) | 0.92 | |

**Supplementary Table S7.** P values comparing FRET latency Δτ ($\tau_{FRET}$ -$\tau_{Ca\ Uptake}$) differences between groups analyzed by 1-way ANOVA with Dunn's *post-hoc* test. See Fig. S8F for summary data.

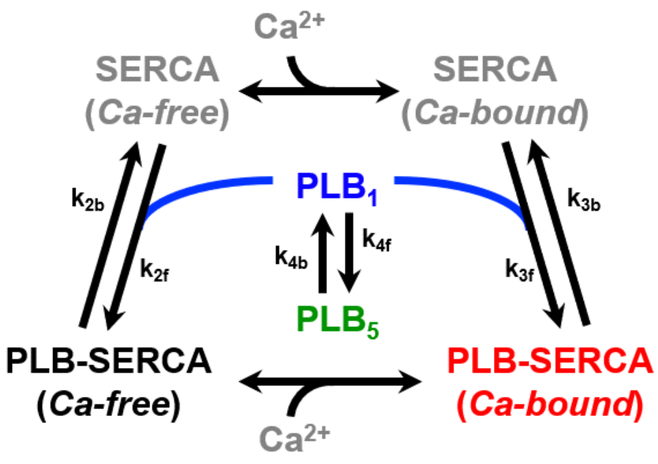


| Parameter | Value ($s^{-1}$) | SE ($s^{-1}$) |
|---|---|---|
| $k_{2f}$ | $8.1 \times 10^3$ | $5.8 \times 10^2$ |
| $k_{2b}$ | $1.1 \times 10^2$ | $3.3 \times 10^0$ |
| $k_{3f}$ | $5.6 \times 10^3$ | $6.7 \times 10^2$ |
| $k_{3b}$ | $1.7 \times 10^2$ | $3.2 \times 10^1$ |
| $k_{4f}$ | $1.0 \times 10^1$ | $9.0 \times 10^{-1}$ |
| $k_{4b}$ | $7.9 \times 10^{-2}$ | $1.8 \times 10^{-2}$ |

**Supplementary Table S8.** Mean kinetic parameters of PLB -SERCA and -pentamer interactions fit from experimental FRET measurements for the computational model.

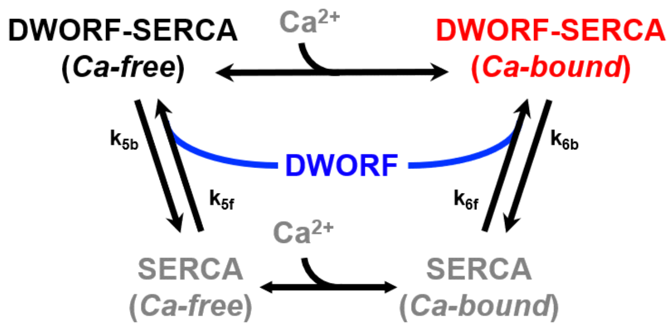


| Parameter | Value ($s^{-1}$) | SE ($s^{-1}$) |
|---|---|---|
| $k_{5f}$ | $3.5 \times 10^1$ | $7.5 \times 10^0$ |
| $k_{5b}$ | $1.9 \times 10^1$ | $8.5 \times 10^0$ |
| $k_{6f}$ | $1.3 \times 10^4$ | $1.4 \times 10^3$ |
| $k_{6b}$ | $4.8 \times 10^3$ | $1.8 \times 10^3$ |

**Supplementary Table S9.** Mean kinetic parameters of DWORF-SERCA interactions fit from experimental FRET measurements for the computational model.

**Supplementary Methods**

To simulate the $Ca^{2+}$ transient, we used the following equations:

$$\mathrm{Ca(t)} = \begin{cases} 0 & t < t_{init} \\ \frac{(t-t_{init})^n}{K+(t-t_{init})^n} & t_{init} < t < t_{init} + t_{setpoint} \\ \frac{K}{K+(t-t_{init})^n} & t > t_{init} + t_{setpoint} \end{cases} \qquad \text{eq. S1}$$

where $t_{init}$ is the time duration of basal $Ca^{2+}$, $t_{setpoint}$ is the time duration of peak $Ca^{2+}$, and $K$ and $n$ are $Ca^{2+}$ transient shape parameters.

As a simplification, we assume that SERCA exists in two ensembles: diastolic and systolic. Furthermore, these two ensembles distribute following the $Ca^{2+}$ transient, which can be described by the following equations:

$$SERCA_{dia,tot} = SERCA_{tot} * \frac{Ca(t)}{Ca_{max}} \qquad \text{eq. S2}$$

$$SERCA_{sys,tot} = SERCA_{tot} - SERCA_{sys} \qquad \text{eq. S3}$$

where $SERCA_{dia,tot}$ is the diastolic SERCA ensemble, $SERCA_{sys,tot}$ is the systolic SERCA ensemble, and $SERCA_{tot}$ is the total amount of SERCA.

The following ordinary differential equations were used to describe the models of PLB-SERCA and DWORF-SERCA binding.

PLB-SERCA

$$\frac{dPLB{\cdot}SERCA_{dia}}{dt} = k_{2f} * PLB_1 * SERCA_{dia} - k_{2b} * PLB \cdot SERCA_{dia} \qquad \text{eq. S4}$$

$$\frac{dPLB{\cdot}SERCA_{sys}}{dt} = k_{3f} * PLB_1 * SERCA_{sys} - k_{3b} * PLB \cdot SERCA_{sys} \qquad \text{eq. S5}$$

$$\frac{dPLB_5}{dt} = k_{4f} * PLB_1 - k_{4b} * PLB_5 \qquad \text{eq. S6}$$

DWF-SERCA

$$\frac{dDWORF{\cdot}SERCA_{dia}}{dt} = k_{5f} * DWORF * SERCA_{dia} - k_{5b} * DWORF \cdot SERCA_{dia} \qquad \text{eq. S7}$$

$$\frac{dDWORF{\cdot}SERCA_{sys}}{dt} = k_{6f} * DWORF * SERCA_{sys} - k_{6b} * DWORF \cdot SERCA_{sys} \qquad \text{eq. S8}$$

where $SERCA_{dia}$ and $SERCA_{sys}$ are free diastolic and systolic SERCA, $PLB{\cdot}SERCA_{dia}$ and $PLB{\cdot}SERCA_{sys}$ are diastolic and systolic PLB-SERCA complexes, $DWORF{\cdot}SERCA_{dia}$ and $DWORF{\cdot}SERCA_{sys}$ are diastolic and systolic DWORF-SERCA complexes, $PLB_1$ is monomeric PLB, $PLB_5$ is pentameric PLB, and $DWORF$ is free DWORF.
